# A microfluidic model of colonocyte-microbiota interaction mimicking the colorectal cancer microenvironment

**DOI:** 10.1101/2023.08.29.555442

**Authors:** Daniel Penarete-Acosta, Rachel Stading, Laura Emerson, Mitchell Horn, Sanjukta Chakraborty, Arum Han, Arul Jayaraman

## Abstract

Changes in the abundance of certain bacterial species within the colorectal microbiota correlate with colorectal cancer development. While carcinogenic mechanisms of single pathogenic bacteria have been characterized *in vitro*, limited tools are available to investigate interactions between pathogenic bacteria and both commensal microbiota and colonocytes in a physiologically relevant tumor microenvironment. To address this, we developed a microfluidic device that can be used to co-culture colonocytes and colorectal microbiota. The device was used to explore the effect of *Fusobacterium nucleatum*, an opportunistic pathogen associated with colorectal cancer development in humans, on colonocyte gene expression and microbiota composition. *F. nucleatum* altered the transcription of genes involved in cytokine production, epithelial-to-mesenchymal transition, and proliferation in colonocytes in a contact-independent manner; however, most of these effects were diminished by the presence of fecal microbiota. Interestingly, *F. nucleatum* significantly altered the abundance of multiple bacterial clades associated with mucosal immune responses and cancer development in the colon. Our results highlight the importance of evaluating the potential carcinogenic activity of pathogens in the context of a commensal microbiota, and the potential to discover novel inter-species microbial interactions in the colorectal cancer microenvironment.

## Introduction

The role of gastrointestinal (GI) tract microbiota in colorectal cancer (CRC) has been extensively investigated^1^, and carcinogenic mechanisms of some bacteria that are abundant in the colorectal microbiota of CRC patients have been identified^2–4^. For example, *in vitro* studies have shown that *Fusobacterium nucleatum* (Fn), an opportunistic pathogen frequently associated with CRC^2^, activates inflammatory and mitogenic transcriptional programs and enhances cell migration in colonocytes upon direct contact^5^. Similarly, colonocytes exposed to a metalloprotease toxin produced by enterotoxigenic *Bacteroides fragilis* display increased cell proliferation, production of reactive oxygen and nitrogen species, and DNA damage^3^. Colibactin, a toxin produced by *Escherichia coli pks+*, induces chromosomal instability and DNA damage *in vitro* and increases tumor formation *in vivo*^4^. Despite these discoveries, the roles of other potential pro-carcinogenic bacteria are not fully understood, which hampers the development of effective CRC prevention and treatment strategies^6,7^.

The interaction between specific bacteria and colonocytes has been primarily studied *in vitro* using co-culture assays; however, these assays fail to capture key elements of bacteria-colonocyte interactions seen *in vivo* in CRC. In these assays, bacteria are added to conventional mammalian cell cultures where they come into direct contact with colonocytes and invade them^6,7^. In contrast, colorectal tumors in the transverse, descending, and sigmoid colon present a thick mucus layer that prevents the pathogenic bacteria from coming in direct contact with colonocytes^8^, thus, these assays ignore potential contact-independent mechanisms of host-microbiota interaction. While most studies focus on interactions between single pathogenic species and colonocytes^6,7,9^, the *in vivo* CRC microenvironment contains hundreds of commensal bacterial species that cross-feed nutrients, compete for niches, or are predatory^10^, and promote colonization resistance against pathogens^11^. Thus, the impact of commensal microbial interspecies interactions on the carcinogenic activity of pathogens in CRC are largely ignored in standard *in vitro* models.

Microfluidic models of healthy GI epithelium such as the gut-on-a-chip^12^ and the HuMiX^13^ devices overcome some of these limitations by enabling the co-culture of intestinal epithelial cells with multi-strain microbial communities, facilitating the study of probiotic-host interaction and inflammatory diseases. While these models have increased our capability to study GI physiology, they do not mimic key pathological features of CRC tissues such as the hypoxic three-dimensional tumor microenvironment experienced by colonocytes *in vivo*^14^ or the anoxia that bacteria encounter in the colorectal lumen^15^, which in turn impact cancer cell metabolism^16^, gene expression^17^, proinflammatory signaling^18^, and response to xenobiotics^14^, as well as bacterial metabolism^19^, and virulence^20^. Therefore, there is a need for more physiologically relevant co-culture models that allows investigation of the interaction between bacteria and colonocytes in a microenvironment that mimics CRC. Here, we report the development of a microfluidic model that facilitates studying interaction between a microbial community and three-dimensionally growing colonocytes under an *in vivo*-like tumor microenvironment. We employ microfluidic perfusion and compartmentalization to control cell localization and oxygen concentrations to sustain viable cultivation of colonocytes in co-culture with a diverse anaerobically cultured microbial community. We utilized this model to investigate the interaction between colonocytes, the pathogen Fn, and commensal microbiota. Our study illustrates the importance of mimicking the CRC microenvironment when studying host-microbiota interaction in CRC.

## Results

### Device design and operation

The microfluidic microbiota-colonocyte co-culture device consists of four stacked microfluidic layers separated by three porous membranes (**Fig. 1A-D**). The middle two layers house the microbiota and colonocytes culture chambers, while the top and bottom channels are used for perfusing appropriate culture media to the respective cell chambers. A 0.2-μm pore size membrane separates the two culture chambers and prevent the migration of bacteria into the mammalian culture chamber. A 0.2-μm pore size membrane also separates the microbiota culture chamber and media flow channel to allow media perfusion without washing out bacteria. The mammalian cell culture chamber and respective culture medium channel are separated by an 8-μm pore size membrane to facilitate diffusion of nutrients and waste removal while keeping the cells within their chamber. Each cell culture channel contains four cell culture chambers that allows replicates to be used for downstream analysis (gene expression, immunohistochemistry, and viability assays).

**Fig. 1:**
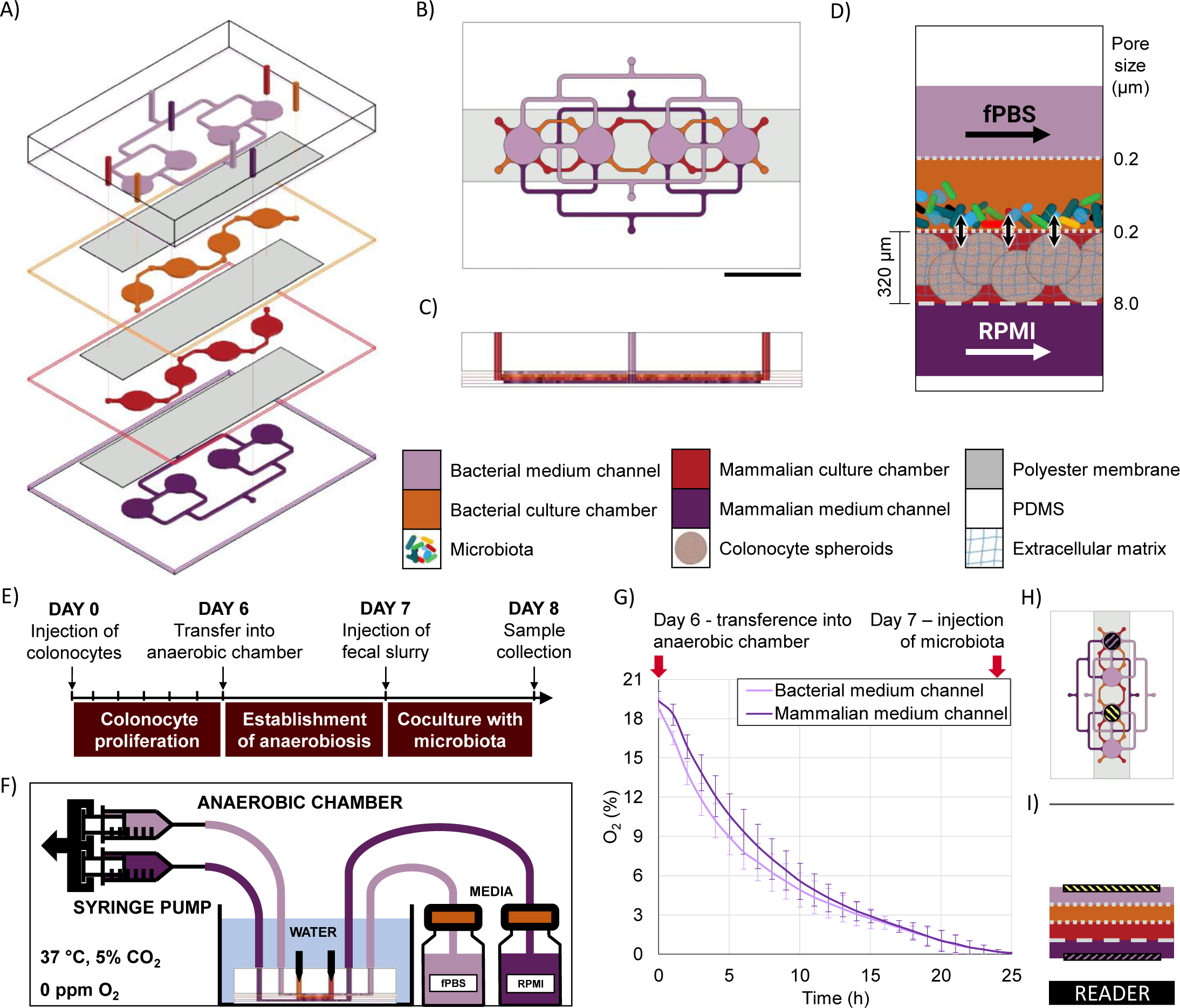
A microfluidic device to study the interaction between bacteria and colonocytes in the colorectal cancer microenvironment. A) Exploded, B) top and C) cross-sectional views highlighting media channels and culture chambers. Scale bar = 1 cm. D) Schematic representation of device cocultures. Confined bacterial and colonocyte populations interact via small molecules during perfusion with growth media. E) Operation schedule. F) Scheme of set-up during operation inside anaerobic chamber, showing device connections to media bottles and syringe pump, as well as conditions inside anaerobic chamber. G) Oxygen quantification inside medium channels during Establishment of Anaerobiosis stage. Error bares represent SEM. H) Top and I) cross-sectional views showing the position of oxygen sensing spots inside the device and the location of the reader.

The device is operated in three stages (**Fig. 1E**). First, a suspension of HCT116 colonocytes in Matrigel is injected into the mammalian cell culture chamber and perfused with RPMI medium under a 21% O_2_ atmosphere for 6 days to allow cell attachment and three-dimensional proliferation. Next, the device is transferred into an anaerobic chamber where the oxygen concentration drops to a negligible level within 24 hours, as confirmed by fluorescence quenching-based oximetry (**Fig. 1F-I**). Finally, a microbial suspension is injected into the bacterial cell culture chamber and co-cultured with the colonocytes for 24 hours. At the end of the co-culture period, the device is disassembled, and cells are harvested for characterization and molecular analysis.

### Effect of culture environment on the expression of cancer-related genes

CRC tumors are characterized by loss of tissue architecture^21^ and heterogeneous oxygen distribution ranging from hypoxia to near anoxia due to poorly developed vasculature^22^. These alterations are associated with changes in the expression of colonocyte genes involved in metabolism, cell signaling, proliferation, inflammation, and epithelial-to-mesenchymal transition (EMT) during cancer^23^. To capture these alterations, we first analyzed colonocytes cultured within a three-dimensional Matrigel scaffold in the microfluidic device. We observed the formation of spheroids (∼ 200 µm diameter) with high cell viability, with well-defined E-cadherin cell-cell adherens junctions (**Fig. 2A-C**). Compared to 2D cultures, HCT116 colonocytes in the device exhibited similar gene expression profiles as spheroids cultured in tissue culture plates. This included genes involved in glucose transport (*GLUT-1*), acid-balance regulation (carbonic anhydrase 9 - *CA IX*), pro-angiogenic and proinflammatory signaling (vascular endothelial growth factor A - *VEGF-A* and interleukin 8 - *IL-8*), control of proliferation (*Ki-67*), cell-cell junction (epithelial cadherin 1 (*CDH1*), cytoskeleton stabilization (*Vimetin*), and the proto-oncogenes *myc* and *KRAS* (**Fig. 2D**). However, the expression of Zinc finger E-box-binding homeobox 1 (*ZEB1)* and *Fibronectin1*, EMT genes that regulate cell migration and cancer dissemination^24^, were decreased while the expression of Zinc finger protein *SNAI2* was increased in 3D-cultured HCT116 cells in the device relative to culture in tissue culture plates.

**Fig. 2:**
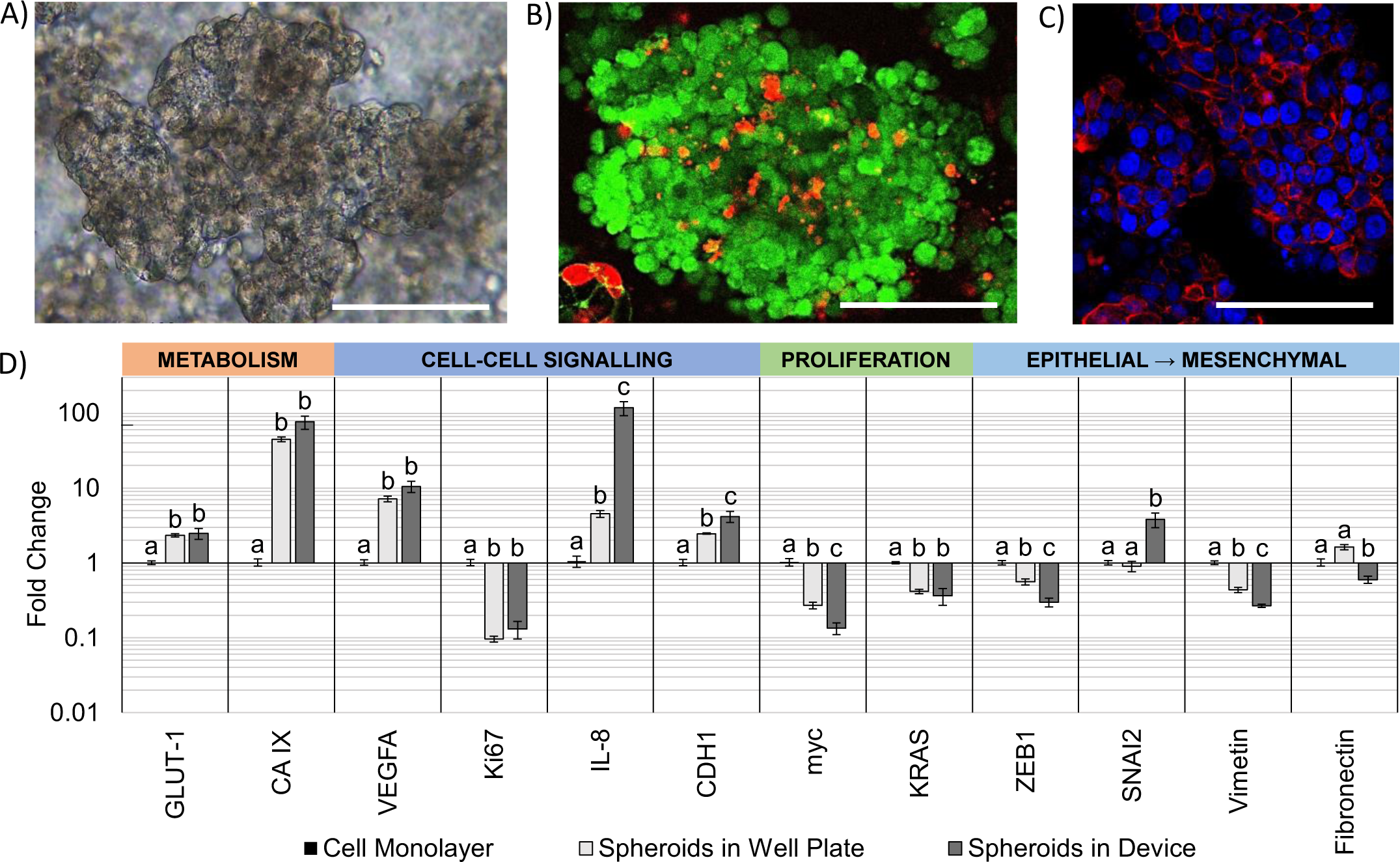
Characterization of cell populations cultured in the microfluidic device. A) Brightfield, B) Live-Dead staining, and E-cadherin (red) staining of HCT116 spheroids grown in the device. Scale bar = 100 µm. D) Changes in gene expression in HCT116 spheroids in the device compared to spheroids in well plates. Different letters indicate statistical significance in difference within each gene (p-value < 0.05, n = 4). Error bars represent SEM.

### Microbial community culture in the microfluidic device

Due to the high diversity of bacterial species in the colon *in vivo*^25^, it is important to sustain a diverse microbial community in *in vitro* models for studying the interaction between colonocytes and the microbiota. However, in vitro culture of microbiota in nutrient-rich media typically results in reduced diversity and increased abundance of members such as proteobacteria^26^. To overcome this challenge, we cultured murine fecal microbiota using PBS supplemented with soluble nutrients present in fecal matter (fPBS). Culturing the microbiota in fPBS resulted in bacterial viability of ∼90%, as well as no decrease in culture turbidity (OD_600_) or colony forming units (CFU) over 24 hours compared to the original inoculum (**Fig. 3A-E**). Metagenomic analysis using 16S rRNA sequencing showed the presence of 40 operational taxonomic units (OTUs) in the fecal inoculum; after 24 hours of culture in the device, 33 OTUs (82.5%) were still detected in the cultured community (**Fig. 3F-G**). Core microbiome analysis, which identifies highly abundant (>1%) and prevalent (>20%) OTUs, revealed that 13 of the 15 members of the core microbiome in the device-cultured community were also present in the core microbiome of the fecal inoculum (**Fig. 3H**), including those characteristic of the murine microbiota such as *Bacteroides*, *Prevotella*, *Mucispirillum*, and the families S24-7, Clostridiales, and Lachnospiraceae. These results demonstrate the feasibility of sustaining a diverse and relevant microbiota community on-chip.

**Fig. 3:**
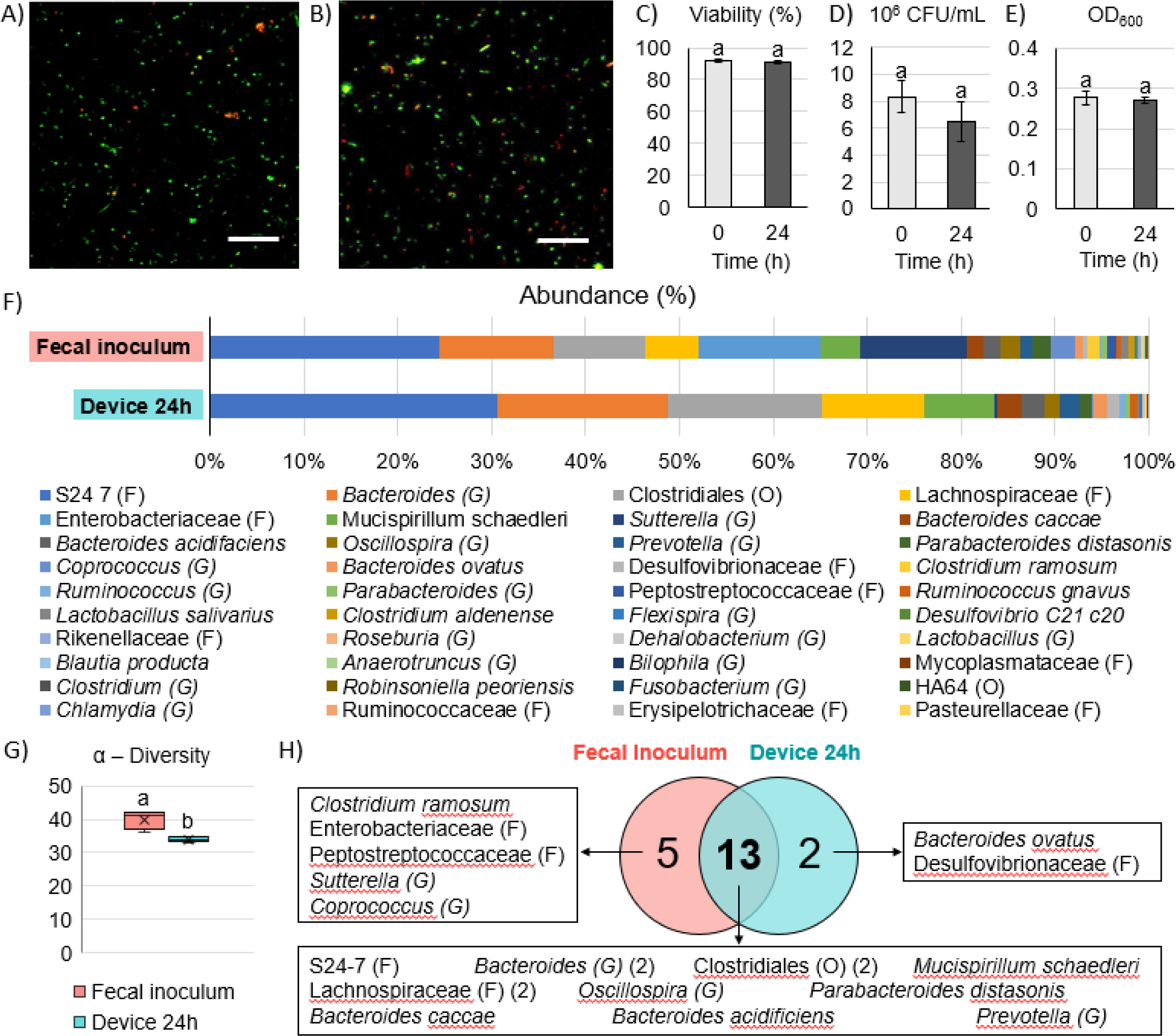
Culture of microbiota in the microfluidic device. A) Live/Dead fluorescent staining of microbiota after injection in device and B) culture for 24 hours with fPBS in the device. Scale bar = 50 µm. C) Changes in viability (Live/Dead staining), colony forming units (CFU), and E) optical density of microbiota after 24 of culture in the device. Letters indicate statistical significance in difference (p-value < 0.05, n = 4). Error bars represent SEM. F) Composition of fecal inoculum and microbiota after 24 hours of culture in the device (n = 4). G) Biodiversity analysis. Letters indicate statistically significant difference (p-value < 0.05, n = 4). H) Core microbiome OTU distribution in communities.

### Co-culture of colonocytes and microbiota in the microfluidic chip

Co-culture of HCT116 colonocytes and fecal microbiota in well plates and transwell inserts in an anaerobic chamber for 24 hours resulted in significant decrease in colonocyte viability (**Fig. 4A/D)** and pH (**Fig. 4E**). This is likely due to the accumulation of bacterial fermentation products and the invasion of colonocytes by bacteria, as previously reported in aerobic cocultures of colonocytes with bacteria^27^. We hypothesized that spatial segregation between bacteria and colonocytes in the microfluidic device, along with pH stabilization by continuous medium flow, would increase the viability of colonocytes during co-culture. Indeed, a significant increase in viability was observed when HCT116 cells were co-cultured with microbiota in the microfluidic device (**Fig. 4B/D**), with approximately 45% of HCT116 cells remaining viable compared to 70% viability in the absence of bacteria (**Fig. 4D**). Stable eluate pH of 7.4 in the mammalian culture channel and 6.8 in the bacterial culture channel were observed, suggesting that continuous removal of acidic bacterial metabolites may have contributed to the improved colonocyte viability. We also confirmed that spatial segregation is necessary to maintain HCT116 colonocyte viability in the device, as removal of the membrane that separates colonocytes from microbiota resulted in bacterial overgrowth and near total loss of colonocyte viability (**Fig. 4C**). Taken together, these results demonstrated that both perfusion culture and spatial segregation in the microfluidic device supports the co-culture of colonocyte spheroids and microbiota.

**Fig. 4:**
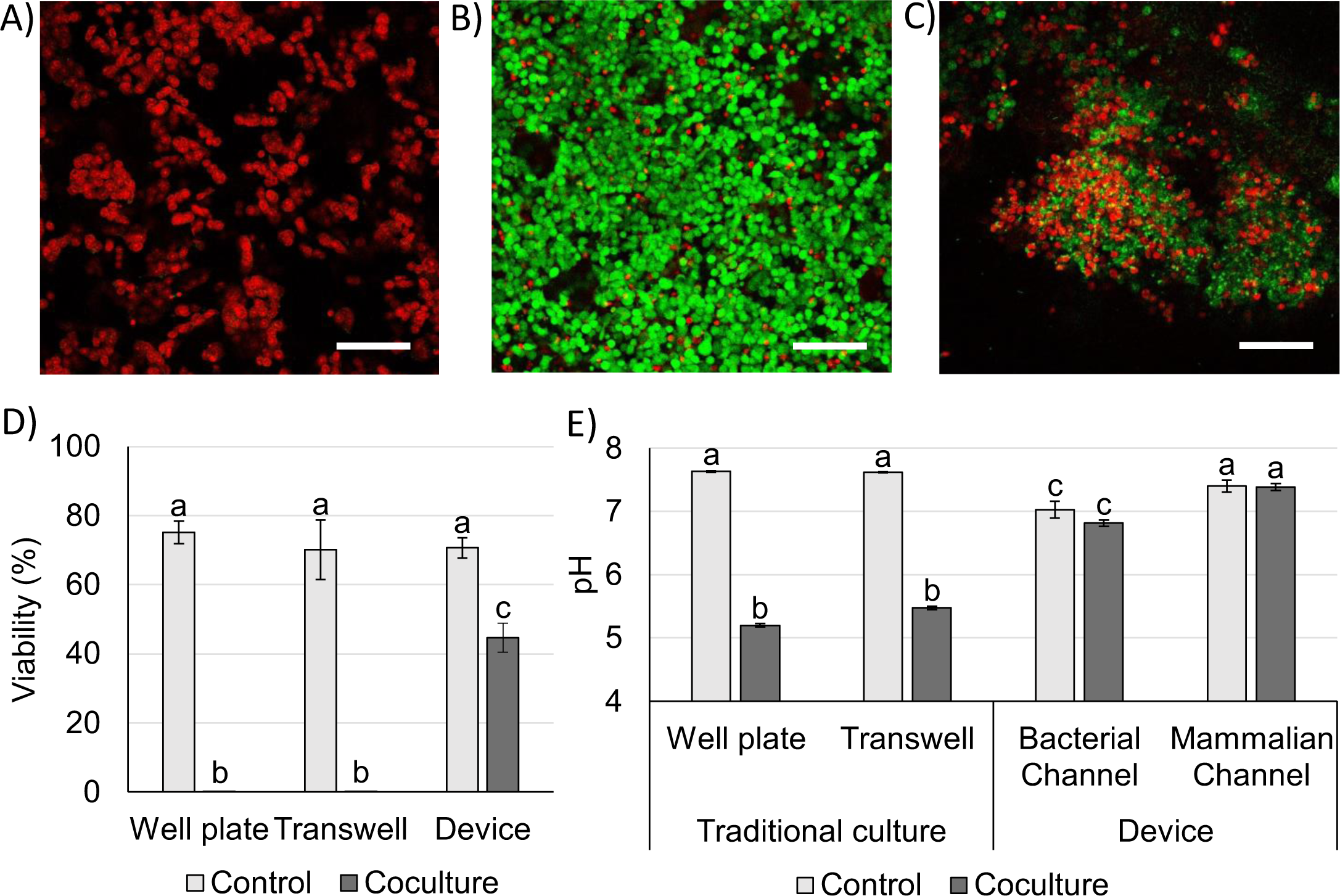
Viability of colonocytes in coculture with microbiota. Representative Live-Dead staining images of HCTT116 cells cocultured with fecal microbiota in A) well plate or transwell insert, B) a microfluidic device, and C) a microfluidic device that permits direct contact between bacteria and colonocytes. Scale bar = 100 µm. D) HCT116 viability and E) pH of culture media in HCT116 controls and cocultures with microbiota in different platforms. Different letters indicate statistically significant differences across all treatments (p-value < 0.05, n = 3). Error bars represent SEM.

### Co-culture of colonocytes with *Fusobacterium nucleatum*

We used the microfluidic co-culture model to explore contact-independent interactions between colonocytes and *Fusobacterium nucleatum* (Fn) (**Fig. 5A**). Co-culture of HCT116 colonocytes with Fn alone resulted in a 2.8-fold upregulation of *IL-8*, a pro-inflammatory cytokine previously reported to be modulated upon direct Fn-colonocyte contact^28^, as well as an upregulation of the matrix metalloprotease gene *MMP1* (3.1-fold) and the EMT genes *Gli* (4.4-fold) and *SNAI2* (8.5-fold), which are linked to metastasis^29,30^ (**Fig. 5B**). Overall, these changes in gene expression are consistent with the reported with Fn using *in vivo* models^31^, demonstrating the capability of the microfluidic device in studying contact-independent interaction between the host and microbiota.

**Fig. 5:**
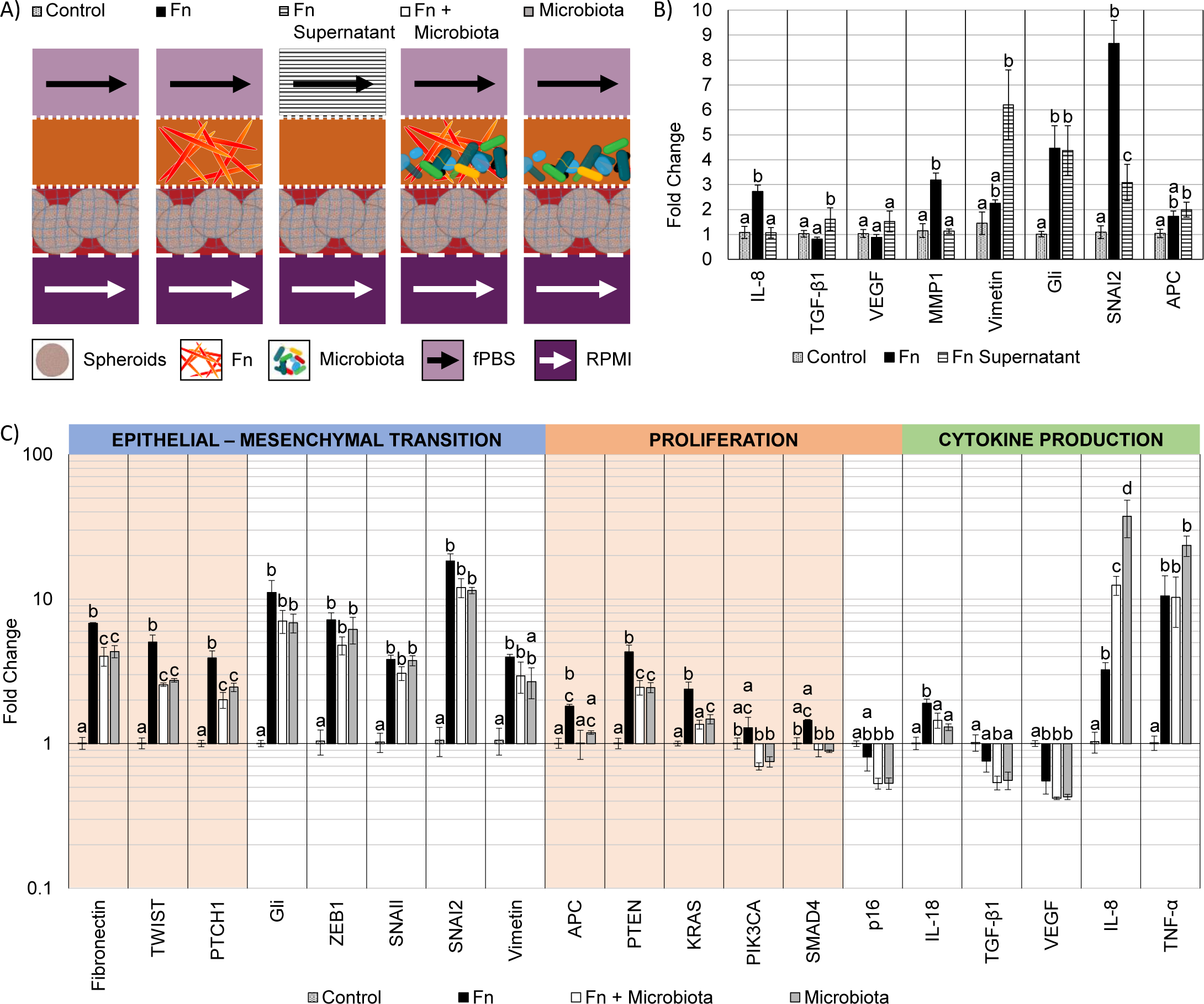
Changes in gene expression in HCT116 upon coculture with microbiota. A) Schematic representation of cocultures in device. B) Fold changes in gene expression of HCT116 upon coculture with Fn or treatment with Fn supernatant, normalized to monoculture control. Letters indicate statistically significant difference within each gene (p-value < 0.05, n = 4). Error bars represent SEM. C) Changes in gene expression in HCT116 induced by Fn in a microbiota. Highlighted genes indicate statistically significant difference between Fn and Fn + Microbiota (p-value < 0.05, n = 3). Error bars represent SEM.

To further understand the changes upon colonocytes and Fn co-culture, we exposed HCT116 colonocytes in the device to cell-free supernatant of Fn cultured in fPBS for 24 hours. Compared to coculture with Fn in the device, exposure of HCT116 colonocytes to the Fn culture supernatant resulted in an upregulation of the EMT genes *Vimentin, Gli* and *SNAI2*, which confirmed that these effects are mediated by contact-independent mechanisms (**Fig. 5A-B**). However, unlike the co-culture with Fn experiment, the Fn culture supernatant had no effect on *IL-*8, *VEGF*, and *MMP1* expression, while *Vimetin* transcription was upregulated 6-fold by Fn supernatant. These results demonstrate the capacity of Fn to induce changes in colonocyte gene expression without direct contact, as well as significant differences between Fn-colonocyte coculture on-chip and treatment of colonocytes with Fn culture supernatant.

### Effect of Fn on HCT116 in the presence of a diverse microbiota

The carcinogenic activity of pathogenic bacteria on colonocytes has traditionally been studied by exposure of colonocytes to a single pathogenic species without considering the combined effect of a diverse microbial community on this interaction. We hypothesized that the presence of a complex microbiota would impact the effect of Fn on HCT116 colonocytes. To test this hypothesis, HCT116 colonocytes were co-cultured in the device for 24 hours with both Fn and fecal microbiota (**Fig. 5A**), followed by colonocyte gene expression analysis. Co-culture of HCT116 with Fn in the presence of microbiota significantly attenuated the gene expression increase observed in co-culture with Fn alone for 8 out of the 19 evaluated genes, including genes associated with EMT and proliferation (**Fig. 5C**). For example, Fn upregulated the expression of the EMT gene *TWIST*, 5-fold, while the presence of the microbiota reduced this Fn-induced upregulation to 2.5-fold. Additionally, 6 genes followed this trend although statistical significance was not determined. Interestingly, addition of Fn to the microbiota decreased the transcription of pro-inflammatory *IL-8* and *TNF-α* that was upregulated by coculture with microbiota. These results demonstrate that a diverse microbiota can attenuate the effect of a pathogen like Fn.

A high Fn abundance has been correlated with overall changes in microbiota composition in CRC patients^8^. Therefore, we hypothesized that Fn could promote changes in the composition of a complex microbial community. To test this hypothesis, we used 16s rRNA sequencing to analyze the composition of the fecal community, with or without Fn, after co-culture with HCT116 colonocytes in the microfluidic device (**Fig. 6A**). The abundance of Fn in these co-cultures was adjusted to represent 63% of read sequences after culture since the abundance of Fn has been reported to be between 25% and 80% in Fn-dominated tumor-associated microbiota in CRC patients^8^. Addition of Fn to the microbiota resulted in the emergence of five previously undetected OTUs, including members of the families Peptostreptococcaceae and Enterobacteriaceae, and changes in the abundance of multiple clades (**Fig. 6B-C**). Interestingly, Fn induced a 40-fold increase in the abundance of the genus Sutterella to 12.6%, and the detection of *Clostridium ramosum,* which was undetectable in the microbiota without Fn, to an abundance of 3.2%. At the same time, Fn decreased the abundance of 10 taxa in the microbiota, including members of the order Clostridiales, the family Lachnospiraceae. Notably, a 0.59-fold decrease in the abundance of *Bacteroides ovatus* was also observed. Overall, while the microbiota attenuated the effect of Fn on HCT116 gene expression of the majority of evaluated genes, Fn simultaneously altered the abundance of multiple proinflammatory taxa. These results demonstrate the possibility of using the developed microfluidic model for studying the interaction between a pathogen and commensal microbiota and the resultant changes in host cell gene expression.

**Fig. 6:**
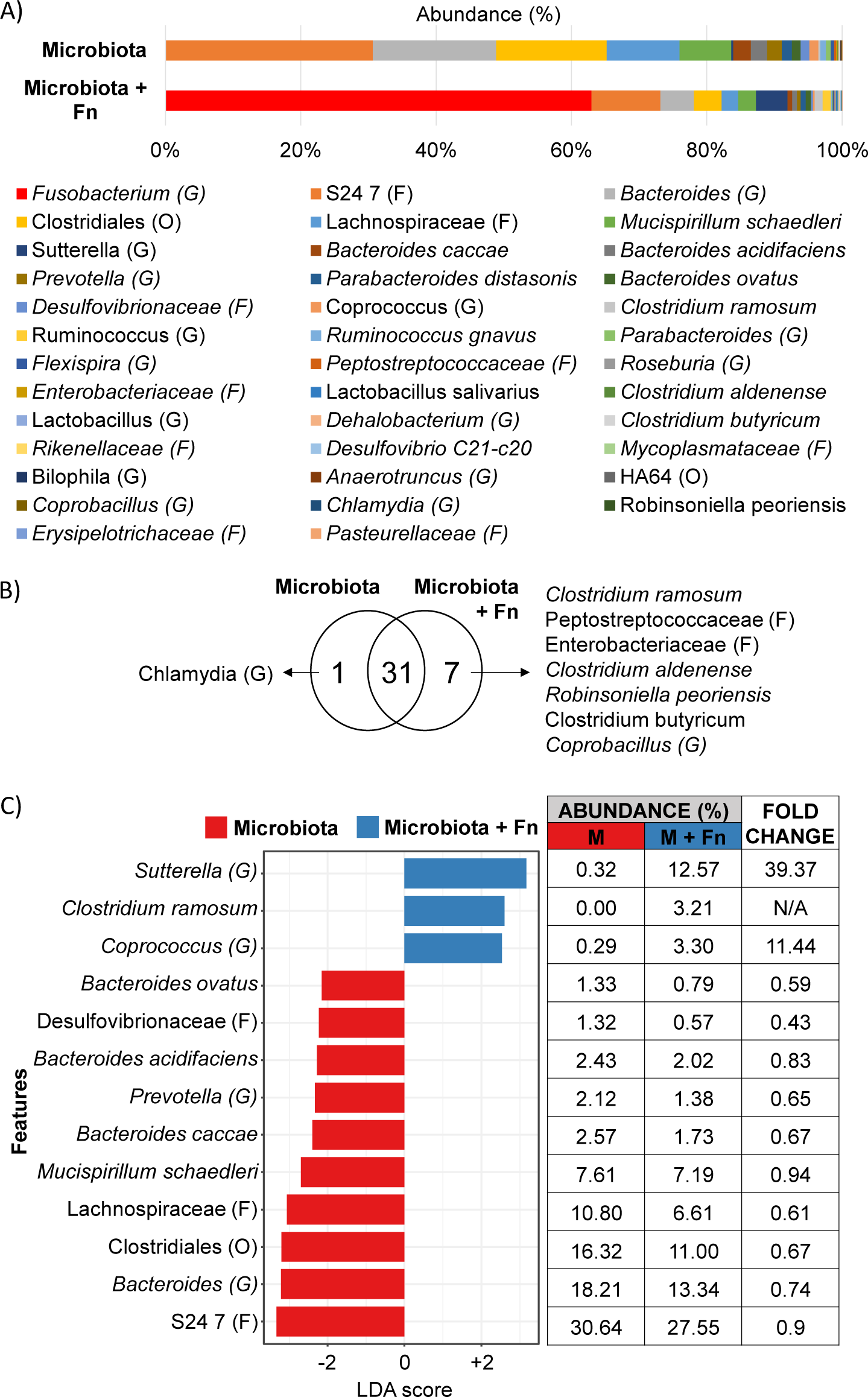
Effect of Fn on microbiota composition during coculture on chip. A) Average composition (n = 4), B) OTU distribution, and C) LEfSe analysis (p-value < 0.05) of microbiota and microbiota with Fn.

## Discussion

While the causal role of microbiota in CRC has been proposed^1^, mechanistic studies on how specific pathogens interact with commensal microbiota and impact host gene expression have been limited. One of the challenges has been that most *in vitro* models used to study colonocyte-microbiota interaction cannot maintain viable colonocytes in the presence of a predominantly anaerobic microbial community. Using a microfluidic device that allows spatial segregation of colonocytes and microbiota by a porous membrane so that they can be co-cultivated while preventing bacterial infection of colonocytes, together with continuous perfusion of growth media that prevents the accumulation of cytotoxic bacterial metabolites, we demonstrate the ability to sustain colonocyte viability in the presence of a microbial community (**Fig. 4**). In parallel, the use of minimal growth medium prepared from feces sustained a diverse microbial community representative of fecal microbiota (**Fig. 3**). Taken together, these design features enabled investigating the interaction between microbiota, specific pathogens and colonocytes in CRC.

Previouslydeveloped microfluidic GI tract models devices have demonstrated co-culturing epithelial cells in a monolayer to form a tight barrier against GI microbiota^12,13^. However, these models of healthy epithelium do not provide an oxygen-deprived 3D microenvironment that is characteristic of tumors^32^. Additionally, these models only provide a hypoxic microenvironment (∼1% O_2_) for the culture of GI microbiota^12,13^, despite the fact that the GI microbiota contains obligate anaerobes^33^. In the microfluidic model described here, colonocytes from the undifferentiated CRC cell line HCT116, embedded in Matrigel (an ECM scaffold known to foster the growth of patient-derived tumor cells into aggregates with preserved adenocarcinoma histology)^34^, grew into spheroids. These spheroids were then co-cultured with fecal microbiota in an anoxic microenvironment that matches the absence of oxygen in the colonic lumen^33^.

Three-dimensional culture of HCT116 colonocytes in our device resulted in marked changes in the transcription of genes related to metabolism, proinflammatory signaling, proliferation, and EMT compared to monolayer cultures in well plates (**Fig. 2D**), in a manner that is consistent with the expected effect of anoxia and 3D culture^17^ and typical of cancerous tissue^32,35^. The most significantly upregulated genes in our model (*GLUT-1*, *CA IX*, *VEGF-A*, and *IL-8*) have been reported to be upregulated in CRC tissues compared to healthy tissue in patients, and are linked to poor prognosis^36–39^. Crucially, the levels of expression of many genes differentially regulated in the device, including *IL-8*, *ZEB1*, *SNAI2,* and *KRAS,* were also significantly impacted upon co-culture with the microbiota (**Fig. 5C**). This highlights the importance of providing a relevant microenvironment to colonocytes during experimental treatments to assess the degree of transcriptional changes induced by the microbiota.

The majority of work on the carcinogenic activity of bacterial pathogens like Fn has focused on the characterization of contact-dependent mechanisms (i.e., effector proteins and intracellular bacterial invasion) of pathogens^5,28^. These mechanisms require invasion of the mucus layer and biofilm formation on the epithelial surface to allow bacteria-colonocyte contact^40^. Such colonization events are common in proximal colon malignancies; however, biofilm formation occurs only in 12% of distal colon tumors^8^. Therefore, contact-independent mechanisms where colonocytes are exposed to the metabolic products of a dysbiotic microbiota dominated by opportunistic pathogens such as Fn are likely important, but not well studied. The changes in expression of cytokines, EMT, and matrix remodeling observed with Fn in a contact-independent manner using our device (**Fig. 5B**), support the likelihood of such mechanisms *in vivo*. To our knowledge, this is the first report of contact-independent effects of Fn on colonocytes in a CRC-relevant microenvironment. These changes in gene expression could be exerted by microbial outer membrane vesicles^41^, metabolites like hydrogen sulfide^42^, and short-chain fatty acids^43^, which require further investigation. It is expected that our model can be used to investigate causality between pathogenic bacteria and transcriptional changes in colonocytes that mimic those observed in CRC tissue.

It was interesting to observe that experiments with cell-free supernatants did not fully capture the contact-independent effects observed with Fn co-culture. Treatment of colonocytes with bacterial culture supernatant has been extensively used to study the carcinogenic activity of bacterial metabolites and virulence factors^44,45^. However, this only represents uni-directional effects between the bacterial supernatant and colonocytes while potentially missing bi-directional effects (e.g., the effect of micro-RNA secreted by the host in extracellular vesicles that potentially target bacterial genes and regulates bacterial metabolism^46^, host-derived glycans influencing the composition of the microbiota by fostering interspecies competition for nutrients^47^). *In vivo*, colonocytes and the microbiota continuously and bidirectionally exchange micro-RNA, hormones, cytokines, and metabolites^46,48^, and thus emulating this type of communication may be key to understanding the role of microbiota in CRC. The observation that the transcription of *IL-8*, a significantly upregulated cytokine in CRC that contributes to tumor growth, invasion, and metastasis^49^, was upregulated only during co-culture with Fn and not after treatment with Fn-free supernatant supports the importance of bi-directional communication in our model (i.e., the sustained production of proinflammatory molecules by Fn in response to colonocyte-derived signaling molecules leads to upregulation of *IL-8*).

The carcinogenic activity of bacterial pathogens has traditionally been evaluated at the single species level, but in the GI tract, hundreds of bacterial species interact^10^ and host cells are exposed to the net product of such interactions^50^. To the best of our knowledge, this is the first *in vitro* study to evaluate the carcinogenic activity of a bacterial pathogen within the context of a complex microbiota. In our model, the presence of a diverse microbiota significantly attenuated the effect of Fn (**Fig. 5C**) even though Fn represented 63% of the OTUs in the community (**Fig. 6**). This suggests that the observed attenuation could be due to the combined effect of metabolites produced by the microbiota and Fn, or by a shift in Fn metabolism due to interspecies competition for substrates such as glycans^47^, micronutrients^51^ or dietary fiber^52^. The gastrointestinal microbiota and its products induce production of pro-inflammatory cytokines IL-8 and TNF-α in mice^53,54^, and this effect is recapitulated in our model (**Fig. 5C**). Interestingly, the presence of Fn in the microbiota significantly reduced the induction of the IL-8 and TNF-α in HCT116 colonocytes, compared to the microbiota alone. This is contrary to the reported pro-inflammatory effect of Fn cultured in direct co-culture with CRC cell lines that has been hypothesized to play a role in Fn-induced carcinogenesis^28^, and needs to be further investigated. This discrepancy again underscores the importance of studying the effect of pathogenic bacteria within a representative microbiota and a relevant microenvironment. Overall, these results demonstrate the usefulness of our model in evaluating pathogen-commensal microbiota interaction and its impact on the host.

It has been hypothesized that CRC-associated dysbiosis may be driven by single species capable of directly remodeling the microbiota or altering the mucosal microenvironment to induce dysbiosis and promote carcinogenesis^55^. This hypothesis has been difficult to test *in vitro* due to the lack of platforms that include a diverse microbiota and the ability to co-culture Fn with a microbiota and colonocytes. Since Fn is a keystone species for biofilm formation in the gastrointestinal tract^56^, we analyzed the effect of Fn on microbiota composition in our CRC model. OTUs belonging to the Families Enterobacteriaceae and Peptostreptococcaceae, which are associated with inflammation and more abundant in CRC patients^57,58^, were detected only upon addition of Fn. Fn also significantly decreased the abundance of short-chain fatty acid producers such as members of the order Clostridiales and the family Lachnospiraceae, which contain species that act as mediators of anti-cancer immune regulators and inhibitors of tumorigenesis, respectively^59,60^. The addition of Fn significantly increased the abundance of the species *C. ramosum* (from undetectable to 3.2%) and the genus *Sutterella* (40-fold to 12.61%), associated with IgA degradation and decreased defense against pathogen colonization^61,62^. Furthermore, the addition of Fn also decreased the abundance of *B. ovatus*, which is reported as an inducer of IgA production in IBD patients^63^. Since secreted IgA inhibits bacterial attachment to and/or invasion of colonocytes^64^, our results suggest that bacterial infection in proximal CRC tumors or distal tumors with compromised mucus barrier could be facilitated by Fn through modulation of the abundance of IgA inducers and degraders. Further studies are required to test this hypothesis. Overall, our results suggest that while a complex microbiota can attenuate the effect of Fn on colonocyte gene expression, Fn can significantly alter the composition of a microbial community, which could contribute to dysbiosis during CRC.

In summary, the microfluidic device described here enabled evaluating the interaction between a carcinogenic pathogen with commensal microbiota and colonocytes in a microenvironment that mimics several key features of CRC tissue. Our results demonstrate the importance of emulating abiotic factors and biological interactions when studying the role of microbiota in cancer and resulted in the identification of novel interactions between a pathogen and members of the gastrointestinal microbiota, which is necessary to realize the potential of microbiome interventions to prevent and treat disease. The possibility of co-culturing different cell types and precisely controlling the culture microenvironment has the potential to facilitate studying complex host-microbiota interactions beyond what has been previously possible. The addition of relevant tumor microenvironment components like stromal or immune cells can further increase the usefulness of microfluidic *in vitro* models. Considering the correlation between pathogenic bacteria, microbial dysbiosis and CRC development, the microfluidic model described here can also facilitate microbiome engineering studies for developing novel therapeutic strategies against CRC^65^.

## Methods

### Mammalian cell culture

The human colorectal cancer cell line HCT116 (CCL-247) was obtained from ATCC and propagated in RPMI 1640 medium (Corning) supplemented with 10% FBS (Atlanta Biologicals), GlutaMAX, HEPES, and NEAA (Gibco). To obtain monolayer cultures, cells were seeded at a density of ∼5x10^4^ cells/cm^2^ on tissue culture-treated petri dishes and incubated at 37 °C in RPMI for 24 hours. For culturing multicellular spheroids, cells were seeded at a density of ∼5x10^4^ cells/cm^2^ in RPMI medium on petri dishes coated with 1.5% (w/v) agarose in water and incubated at 37 °C in RPMI for 4 days.

### Preparation of fecal PBS

Freshly-voided fecal pellets were obtained from 6 to 8 weeks-old wild-type C57BL/6 female mice fed a standard chow diet. Feces were collected aseptically into anaerobic jars (BD Port-A-Cul™) and dissolved in anoxic PBS inside an anaerobic chamber, to a final concentration of 250 mg/mL. This slurry was filtered through a 20 µm strainer, which resulted in a filtrate containing bacteria, small undigested food particles, and soluble molecules, and a retained cake containing large undigested food particles. The filtrate was centrifuged at 2000g for 10 minutes, resulting in a pellet enriched in bacterial cells and small food particles, and a supernatant containing soluble nutrients. The supernatant was collected, and the pellet was resuspended in anoxic PBS (final OD_600_ = 30) and used as inoculum for device and batch coculture experiments. The filter cake was homogenized using stainless steel beads (Precellys®, 5500 RPM, 20 seconds, one bead per tube), and resuspended in the previously collected supernatant to harvest some of the nutrients available in the cake. The resulting mix was centrifuged for 1 min at 2000g, and the supernatant filtered through a 0.45 µm filter and diluted 20-fold in PBS, which resulted in a solution that is hereafter referred to as fecal PBS (fPBS).

### Fusobacterium nucleatum culture

*F. nucleatum* subsp. nucleatum Knorr (25586) (Fn) was obtained from ATCC and propagated from frozen stock on BD BBL™ Brucella Agar plates with 5% Sheep Blood, Hemin and Vitamin K_1_ (BD Biosciences) in an anaerobic chamber (Coy Laboratories). Anoxic RPMI for liquid cultures was prepared by supplementing RPMI with 0.5 g/L cysteine (Alfa Aesar), 0.5g/L of hemin (Sigma-Aldrich) and 0.2 mg/L of Vitamin K (Spectrum Chemical) and allowing it to become anoxic inside an anaerobic chamber overnight. For device experiments, 25 mL of anoxic RPMI were inoculated with a single colony of Fn and incubated at 37 °C and 5% CO_2_ under anaerobiosis for 24 hours. Cells were harvested by centrifugation at 3000g for 10 minutes for injection into the device (see below). For supernatant experiments, fPBS was inoculated with Fn (∼8.7x10^5^ CFU/mL) and incubated at 37°C under anaerobiosis for 24 hours. After incubation, cells were centrifuged at 3000g for 10 minutes, and the supernatant was filtered through a 0.45 um syringe filter, and immediately use. For CFU/mL determination, serial dilutions of bacterial cultures were plated in Brucella Agar in triplicate and incubated for 24 hours at 37°C under anaerobiosis.

### Batch coculture experiments

For batch cocultures of colonocytes and a fecal microbiota, 5x 10^4^ HCT116 cells in 0.75 mL of RPMI medium were seeded into petri dishes and incubated at 37 °C overnight under normoxia followed by 24 hours of anoxia. Then, 0.75 mL of fPBS containing 1 μL of fecal bacteria slurry (OD_600_ = 30) were added to the culture and incubated for 24 hours. Indirect cocultures using transwell plates were performed by seeding ∼4.4x10^4^ HCT116 cells in 1 mL of media per well, and 0.5 mL of RPMI medium per insert. The cultures were incubated at 37 °C under normoxia overnight followed by 24 hours of incubation under anoxia. Then, media was replaced with a 1:1 mixture of anoxic RPMI and fPBS, 1 μL of fecal bacteria slurry (OD_600_=30) were added to the to the insert, and coculture continued for 24 hours. The ratio of colonocytes to fecal bacteria during these experiments was set to match the ratio from coculture experiments using the microfluidic device.

### Microfluidic device design and construction

A microfluidic device was developed to coculture colonocytes and a fecal bacterial community. The device consists of four PDMS layers separated by three porous polyester membranes (**Fig. 1**). For building the device, thin patterned PDMS layers were obtained by pouring uncured PDMS mix (Sylgard 184®) on a 3D-printed patterned mold (Stratasys, Inc.) with a total height of 500 μm and a pattern height of 160 μm. Excess uncured PDMS was removed from the top of the mold by pressing down with a heavy block, and then the PDMS was cured for 1 hour at 70 °C. The thin PDMS layers were then peeled off from the mold, and biopsy punches were used to create culture chambers (D = 5 mm, final chamber volume of ∼10 μL) and open access to the perfusion channels and culture chambers (D = 1 mm). The device was assembled layer-by-layer using a thin layer of uncured PDMS as glue, and a polyester membrane was sandwiched between each pair of layers. The construct was cured for 15 minutes at 70 °C after adding each new layer. For tubing connection, a 5-mm thick PDMS lid was perforated at the inlet coordinates for the channels and chambers by using a 1-mm diameter biopsy punch. Then, the top and bottom layers in the stack were sealed with the perforated PDMS lid and a 1-mm thick unpatterned PDMS layer, respectively, by oxygen plasma treatment (5 minutes, 30 W, Harrick Plasma). Finally, the bottom of the device was attached to a glass slide using uncured PDMS that was then cured by baking at 70 °C for 1 hour.

PDMS-filled 10 μL micropipette tips were used as stoppers for the inoculation ports to the mammalian and bacterial culture chambers. The device was connected to media reservoirs by flexible 23-Gauge Tygon® medical tubing (Saint Gobain) using 20-Gauge stainless steel connectors detached from dispensing needles (Jensen Tools). The relatively large dimensions of the channels and low flowrate resulted in a low internal pressure compared to photolithography devices with smaller features, leading to bubble formation during operation. To minimize the formation of bubbles, the assembled device was placed underwater and vacuumed to a final pressure of 3x10^-2^ mbar for 24 hours; then, the device was autoclaved (121 °C, 16 PSIG, 45 minutes) and kept covered in sterile water during operation. This protocol sterilized the device and prevented bubble formation. The device was held underwater for the duration of the experiment and only brought out of water for cell injection.

### Oxygen measurement experiments

To confirm the formation of an anoxic environment inside the coculture chambers in the device, a fluorescence-quenching oxygen measurement system was employed (Scientific Bioprocessing). Oxygen sensor spots were attached to the roof and bottom of the bacterial and mammalian medium channels using PDMS glue. The device was then placed on an oxygen reader to continuously monitor oxygen tension. For this experiments, all three membranes in the devices had an 8-um pore size, as membranes with this pore size were translucid and therefore maximized fluorescent signal quality. This selection allowed us to read oxygen tensions in the upper channel from the bottom of the device through three polyester membranes (**Fig. 1 G-I**). The devices were brought into the anaerobic chamber and covered in anoxic water, and the changes in oxygen tension inside the chip were followed for 24 hours.

### Device operation

For colonocyte injection, sterilized devices were transferred into a laminar flow cabinet and a cell suspension of 5x10^6^ cells/mL in a 50% v/v Matrigel diluted in RPMI medium was injected into the mammalian culture chamber, which polymerized into a hydrogel after 15 minutes of incubation at 37 °C. To allow colonocyte handling recovery and aggregate formation, the device was incubated for 6 days at 37 °C in a normoxic 5% CO_2_ incubator and perfused at a rate of 1 μL/min with antibiotics-free RPMI through the mammalian medium channel and PBS through the bacterial medium channel using a syringe pump (Chemyx). On day 6, the device was transferred into an anaerobic chamber and covered with anoxic water (0.5 g/L cysteine, water column height: 1 cm). Both channels were perfused at a rate of 1 μL/min, and the device was incubated at 37 °C for 24 hours to allow the oxygen to diffuse out of the PDMS. On day 7, the devices were injected with either fecal slurry (OD_600_ = 30), a suspension of *F. nucleatum* (∼3.2x10^8^ CFU/mL resuspended in either anoxic RPMI or fecal bacteria slurry), or sterile anoxic RPMI as control. The coculture proceeded for 24 hours inside the anaerobic chamber under perfusion at 10 μL/min, with fPBS flowing through the bacterial medium channel and RPMI flowing through the mammalian culture channel. For sample collection, the devices were taken out of the anaerobic chamber, and bacterial cells were collected by pipetting. The devices were carefully disassembled by peeling off the layers to collect the mammalian hydrogels.

### Viability evaluation and immunofluorescence

HCT116 cell viability upon coculture was evaluated by fluorescent Live/Dead staining. Whole hydrogels were stained by incubation in PBS containing 2 μM Calcein AM (Enzo Life Sciences) and 1.5 μM Propridium Iodine (Invitrogen) for 15 minutes at 37 °C, and fluorescence was captured using a Leica TCS SP5 confocal microscope. For viability quantification, stained hydrogels were placed on ice in PBS with 10 mM EDTA and then disaggregated by pipetting to obtain single cell suspensions. Bacterial cell viability was evaluated by fluorescent Live/Dead staining (Filmtracer™ Live/dead™ Biofilm Viability Kit, Invitrogen), following manufacturer’s instructions. Fluorescent images were processed using the Analyze Particles function in ImageJ.

For immunofluorescence, collected hydrogels from the device were stained with Human/Mouse E-Cadherin Antibody (R&D Systems) at a final concentration of 5 µg/mL, followed by Donkey Anti-Goat IgG NorthernLights™ NL557-conjugated Antibody (R&D Systems), according to manufacturer’s protocols. Fluorescence images were captured using a Leica TCS SP5 confocal microscope.

### Gene expression analysis

RNA was extracted from HCT116 cells using the RNeasy Mini Kit (QIAGEN) following the manufacturer’s instructions. Genomic DNA in the extracted RNA was eliminated by digesting with DNAse (QIAGEN). cDNA synthesis was performed using qScript™ cDNA SuperMix (QuantaBio) using 100 ng of RNA sample in a 10 µL reaction. Quantitative PCR was carried out in a Lightcycler® 96 (Roche) using FastStart Universal SYBR Green Master (Roche). Primers were designed using Primer Blast (NCBI), and amplicon size and specificity were confirmed by melting peak analysis and agarose gel electrophoresis of the reaction products. Each reaction mix contained 1/40^th^ of the cDNA pool obtained per sample and a total primer concentration of 400 nM. The PCR regime consisted of preincubation for 10 minutes at 95 °C and 45 amplification cycles (95 °C x 15 s, 65 °C x 30 s, 72 °C x 45 s). Data were processed using the 2^-ΔΔCt^ method. Multiple genes were evaluated as endogenous qPCR controls, including *18s*, *YWHAZ*, *PMM1*, *UBC*, *IPO8*, and *VPS29*; from these genes, *UBC* showed the most stable expression level and was employed as endogenous control. Sequences for all used primers are provided in **Supplementary Table 1**.

### 16S rRNA sequencing and bioinformatic analysis

DNA from bacterial communities was extracted by using the DNeasy PowerSoil Kit (QIAGEN) according to manufacturer’s instructions, and sequencing of the v4 region of the 16S rRNA gene was performed (Microbiome Insights). FASTQ files were imported into QIIME2, denoised, and aligned to the Greengenes database. OTU abundance data was then imported into Microbiome Analyst, singletons were discarder, and low count features were removed based on a minimum count of 4 and a prevalence lower than 50% across all samples. Data was scaled using Cumulative Sum Scaling. After these processing steps, the resulting pool of samples contained a total of 40 OTUs at taxonomic levels ranging from Order to Species. Biodiversity, Core Microbiome (20% sample prevalence, 1% relative abundance), and LEfSe analysis were performed at the feature level using Microbiome Analyst. To assess the effect of Fn in the composition of the microbiota, Fn counts were removed from the total counts for all samples, and the abundance of the rest of the community members was normalized to 100% for statistical analysis.

### Statistical analysis

For testing statistical significance, unpaired Student’s t-tests were performed on sets of data with two experimental conditions. One-way ANOVAs were used for comparisons among multiple experimental conditions and during qRT-PCR data analysis. For qRT-PCR data analysis, significance in gene expression changes was determined by comparing ΔC_t_ values across treatments, as gene expression data is log normally distributed^66^. The assumption of equality of variances among data sets was confirmed by using the Levene’s test, and normality was validated using the Shapiro-Wilk test. All experiments were performed in triplicate.

## Acknowledgements

We would like to acknowledge Scientific Bioprocessing, Inc. for their collaboration with the oxygen measurement experiments employing the ID Developer kit. This work was partially supported by funds from the Ray B. Nesbitt Chair endowment to A.J.

## Author Contributions

D.P. and A.J. designed the research. D.P., R.S., L.E. and M.H. performed the experiments. D.P., S.C., A.H. and A.J. analyzed the data. D.P. and A.J. wrote the article with input from A.H and S. C. All authors reviewed, discussed, and edited the manuscript.

