## Supplemental Table 1 for "A microfluidic model of colonocyte-microbiota interaction mimicking the colorectal cancer microenvironment"

### Supplementary material

Supplementary Table 1 Primer sequences for qRT-PCR analysis.

| Gene | Forward Primer Sequence | Reverse Primer Sequence |
|---|---|---|
| APC | TTGGCTCGATGCTGTTCCCA | AGCCCAGGGAGGCGTACATA |
| CA IX | GTTCACCTCAGCACCGCCTT | GCACTGTTTTCTTCCGGGCC |
| E-cad | ACGTACAAGGGTCAGGTGCC | TGGGGTATTGGGGGCATCAG |
| Fibronectin | ACAACACCGAGGTGACTGAGAC | GGACACAACGATGCTTCCTGAG |
| Gli | ACAGAAGGACTGTCTGGCCC | CAGTGCCCGCTTCTTGGTCA |
| GLUT-1 | TACCGCCAGCCCATCCTCAT | CCTGCTCGCTCCACCACAAA |
| IL-18 | GTGTTCAAGACCAGCCTGACCAAC | TAGCTGGGATTGAGGGCATGCG |
| IL-8 | TGGACCACACTGCGCCAACA | TCTCCACAACCCTCTGCACCCA |
| Ki-67 | TGTCACCACCCACAGCCACC | GTTGTTCCTGCCACCGTGCC |
| KRAS | CCAGGCCCTGTGTGAACCTT | ACACTGGATGACCGTGGGGA |
| myc | ACAACACCCGAGCAAGGACG | GCAGCTGCAAGGAGAGCCTT |
| p16 | AGTTACGGTCGGAGGCCGAT | CCACCAGCGTGTCCAGGAAG |
| PIK3CA | AGCGTTTCTGCTTTGGGACAA | CTGATGATGGTCGTGGAGGCA |
| PTCHD1 | TGAGACTGACCACGGCCTG | ACCCTCAGTTGGAGCTGCTTG |
| PTEN | AACGATGGCTGTGGTTGCCA | GCACAGTTCCACCCCTTCCA |
| SMAD4 | TCGACAGATGCAGCAGCAGG | GGATGTTTCCTGCCACGGCT |
| SNAI2 | TCCATTCCACGCCCAGCTAC | GGACTCACTCGCCCCAAAGA |
| SNAIl | ACTATGCCGCGCTCTTTCCT | CGTAGGGCTGCTGGAAGGTA |
| TGF-β1 | GAGTTGTGCGGCAGTGGTTGAG | ACCCGTTGATGTCCACTTGCAGT |
| TNF-α | GCCCAGGCAGTCAGATCATC | CTCTGATGGCACCACCAGCT |
| TWIST | GCGGCCAGGTACATCGACTT | GCTGCAGCTTGCCATCTTGG |
| UBC | GCCGGGATTTGGGTCGCAG | CACGAAGATCTGCATTGTCAAGTG |
| VEGFA | GCGACAGGGGCAAAGTGAGT | GACTGGTCAGCTGCGGGATC |
| Vimetin | TGCGCCTCCGGGAGAAATTG | ACGTGCCAGAGACGCATTGT |
| ZEB1 | AGCGCTTCTCACACTCTGGG | TGCTGTCACGTTCTTCCGCT |
